# The Effects of Teaching Emotion Words on Emotional Granularity, Emotion Regulation/Dysregulation, and Satisfaction with Life

**DOI:** 10.64898/2026.08.12.744382

**Authors:** Jennifer M.B. Fugate, Samuel Kalmus, Helenna Shcherbinin, Molly McKillip, Whitney Shae, Jayden Kasiska-Pettersen

## Abstract

Emotional granularity (EG) has been associated with adaptive emotion regulation, psychological well-being, and reduced psychopathology. The present study examined whether teaching highly specific emotion words would increase EG and improve emotion regulation, emotional dysregulation, and well-being outcomes. A total of 95 adults were randomly assigned to either a 14-day emotion word training condition (n = 48) or a matched control word training condition (n = 47). Before and after training, participants completed assessments of word knowledge, self-reported and behavioral estimates of EG, emotion regulation, emotional dysregulation, and satisfaction with life. Mediation analyses evaluated whether changes in EG explained the relationship between increases in word knowledge variables and positive psychological outcomes. Participants in both emotion and control word conditions showed improvements in word knowledge variables after training, but only those in the emotion word group showed that increased emotion word usage was associated with higher EG (using the Range and Differentiation of Emotion Expression Scale, RDEES) and lower emotional dysregulation and reduced suppression. Those in the control group showed mixed effects for self-reported EG and outcomes. For neither group, however, were any of the effects mediated by either EG measurement. Furthermore, behavioral and self-report measures of EG were not significantly correlated, suggesting that they may capture distinct aspects of emotional functioning. These findings provide preliminary support for emotion vocabulary training as a low-cost intervention that may enhance emotional functioning and suggest that self-perceived EG may be more closely linked to adaptive outcomes.

---

A growing body of evidence links early emotion vocabulary to the development of emotion regulation, suggesting that a child’s understanding of emotion words may directly enable them to regulate emotions more effectively [1–4]. Emotion word knowledge is theorized to provide access to emotion concepts that organize affective sensations into meaningful emotional experiences, thereby transforming diffuse affect into differentiated emotional states [5] (for consistent neural evidence, see [6]). From a constructionist perspective, emotion words function not merely as labels but as conceptual tools that support the learning and application of emotion categories [7–9]. Consequently, training in emotional language may enhance the brain’s predictive capacity, enabling more precise emotion construction and, in turn, more effective emotion regulation [9–12].

The study of alexithymia (a disorder of affect dysregulation) also provides indirect evidence supporting the link between emotional language and emotional regulation [13–14]. Individuals with higher levels of alexithymia appear to produce fewer emotion words and synonyms for target emotion words than those with lower levels of alexithymia [15]. In another study, emotional awareness mediated the relationship between alexithymia and emotion regulation [16]. Adolescents’ baseline alexithymia levels correlate with emotion regulation, such that low levels predict future regulation difficulties [17], and the relationship between alexithymia and psychological distress is mediated by emotion regulation in adults [18]. Together, these studies suggest that individuals with high levels of alexithymia experience limitations when expressing and processing emotional language, which can affect their ability to regulate emotions.

## Emotional Granularity

Individuals show incredible variation in emotion word knowledge and the ability to communicate emotional experiences precisely [19]. This individual difference is termed emotional granularity (EG) - the degree to which individuals represent and label emotional experiences in a differentiated manner [19–22]. For instance, individuals with low EG might not be able to distinguish between “anger” and “frustration”, whereas those with higher EG can distinguish between feelings of “irritation”, “impatience”, “agitation”, etc.

A substantial body of research now supports the idea that individuals who exhibit high EG have better well-being outcomes [21, 23–25] and suffer less from various psychopathologies (e.g., major depressive disorder, social anxiety disorder, autism, eating disorders, self-injury, and borderline personality disorders) [26–30]. Furthermore, increased EG is related to reduced symptoms of anxiety and depression in the general population and is thought to be a transdiagnostic vulnerability across a range of mental health disorders [21]. High EG can also help to protect individuals from maladaptive behaviors or negative outcomes. For instance, individuals with high EG are less likely to engage in self-harm [31], are less prone to maladaptive behaviors (e.g., binge eating and alcohol abuse) [32–33] and impulsive and aggressive behaviors [34–35]. In addition, individuals who self-reported a mental health diagnosis (compared to those who did not) have lower EG for emotion categories [36].

Emotional granularity is also thought to support healthy emotional regulation [37–40]. For example, individuals with low EG have an early reactivity to affective stimuli, which contributes to disadvantages in the appropriate regulation of their emotions [41]. Additional evidence supports this idea: low EG is associated with ineffective use of emotion regulation strategies during periods of high emotional intensity (e.g., exam taking) [42, Study 2]. Specifically, one recent model [16] suggests that EG supports emotion regulation because the increased ability to assign meaning to emotional experiences increases subtle distinctions between emotions that, in turn, enable effective regulation strategies. Finally, individuals with higher EG (especially for negative emotions) reported improved psychotherapy responses for depression and stress, although not for anxiety disorders [43]. Together, the research shows that individuals with high EG have more beneficial outcomes and better emotion regulation than individuals with less differentiated emotions.

## Emotional Granularity Measurement

Emotional granularity can be operationalized in distinct ways, reflecting either momentary emotion labeling processes or trait-like beliefs about emotional differentiation. Accordingly, researchers have employed both behavioral and self-report measures to capture different aspects of the construct. Emotional granularity is typically assessed using experience sampling methods (ESMs), in which participants use a handheld device (e.g., smartphone, tablet) to respond to prompts about their feelings (e.g., levels of endorsement of emotion words used to describe their current feelings). ESM leverages the intensity and variability of “real-world” emotional phenomena and is less subject to retrospective memory biases than survey methods [44]. Emotional granularity is established by calculating the interclass correlation (ICC) between similarly-valenced emotion words endorsed by the participant at either a particular time-point, a daily level, or across “waves” (i.e., treatments) [32, 45]. The ICC represents the degree to which emotion terms are used interchangeably across instances, such that a high ICC reflects emotion term endorsement that lacks precision and is therefore considered low in granularity. Lab-based emotion perception tasks, which similarly calculate ICCs in response to emotional scenarios or videos or vignettes (e.g., Scenario Rating Task, SRT) [46], are also used to compute EG [45].

Survey-based measures (e.g., the Range and Differentiation of Emotion Experience, RDEES [47], and the Toronto Alexithymia Scale, TAS [48] have been used as self-reported, global measures that emphasize trait-based qualities thought to be related to EG and similar constructs of emotional differentiation [49]. In one study, higher emotion word accuracy (i.e., ability to pick the correct definition of an emotion word from related emotion definitions) and self-reported understanding of emotion words resulted in less emotional dysregulation after accounting for self-reported EG (using the RDEES) [50]. Therefore, whether self-reported measures of EG overlap with behaviorally-based measurements (obtained from ICCs of emotion endorsement) is still a matter of debate [51].

Some studies have shown that EG is not associated with emotional awareness but is associated with emotional clarity [52–53]. Other studies have discovered that EG for negative emotions (but not for positive) is associated with both awareness (using the “awareness” subscale of the Trait Meta-Mood Scale; TMMS) [54] and with clarity (using the “difficulty in identifying feelings” subscale of the Toronto Alexithymia Scale; TAS) [48]. While more recent studies view both EG and emotion clarity as related, non-fixed traits, it is unclear how the two constructs might relate or overlap [55], calling into question the extent of the EG nomological net.

## Enhancing Emotional Granularity

Despite the known role of EG in overall mental health and well-being, few studies have attempted to specifically increase EG. In one study, [36] participants who increased emotion concept knowledge (compared to a control group who increased their knowledge about countries) increased the level of negative (but not positive) granularity, with changes persisting over time. While other studies using mindfulness-based interventions report increased EG, they do not explicitly aim to enhance or manipulate EG [56–57].

## Summary and Hypotheses

The literature suggests that emotion words can help direct attention to commonalities across disparate instances, thereby helping people to organize emotion concepts. Such commonalities can serve as cues for identifying one’s emotional state and guide appropriate responses in new situations, thus facilitating emotional regulation [7–8, 39, 58]. Emotional granularity is related to better emotional regulation [38–40]. Therefore, emotion words might support EG, representing a critical step in how an individual’s emotion-word knowledge supports their emotional regulation.

The approach for the current study was to train a treatment group to increase their emotion word knowledge by providing a rich source of visual, in-app training material on highly precise emotion words over two weeks. Participants randomly assigned to a control group learned length- and difficulty-matched control words in the same way over the same period. We measured participants’ self-reported ability to understand, use, and accurately pick the correct definition of each emotion word and control word at baseline and after the two-week training period. We computed EG using both traditional emotion-word endorsement of current feelings (through ESM methods across a maximum of 70 occurrences) and self-reported measures (TMMS, RDEES), which were collected at baseline (pre) and again after the training period (post). We also recorded participants’ emotional regulation strategy use (Emotional Regulation Questionnaire; ERQ) [59], emotional dysregulation (Difficulties in Emotional Reactivity Scale; DERS) [60], and overall well-being (e.g., satisfaction with life; SWL) [61] and mental health continuum; MHC-SF) [62]), at both timepoints. Participants assigned to the emotion word training condition were expected to show greater increases in EG than participants assigned to the control word condition. We further hypothesized that increases in EG would mediate the relationship between emotion word learning and (a) reduced emotional dysregulation, (b) reduced use of expressive suppression, and (c) improved well-being. A secondary aim of this study was to address whether self-reported survey measures of EG correlated with behavior-based measures of EG.

## Methods

### Participants

We used [63] estimates of mediated effects to determine an approximate sample size based on 80% power, using a bias-corrected bootstrap test. Based on previous studies’ effect sizes, we assumed that the alpha path was large for all outcomes (approximately 0.59) and the beta path effect was between small-medium (approximately .26) and medium (approximately 0.39). Using the smaller beta effect, the sample size estimate was 115; for a medium-sized effect, the estimate was 54. Therefore, we aimed to enroll 100 individuals to achieve acceptable power.

We recruited 107 University [(removed for identity)] affiliates (graduate students, faculty, and staff) and community members in an urban area to participate in this study. Participants needed to be 18 years or older and English-speaking from birth. This study was approved by [(University removed for identity)] IRB 1942774. Participants received up to $100 in gift cards over the course of their participation (i.e., session 1, $10 USD; pre-intervention ESM period, $10 USD; word intervention ESM period, $50 USD; session 2, $20 USD, post-intervention ESM period, $10 USD). Table 1 provides the participant demographics. There were a total of 48 in the emotion word group and 47 in the control word group after data cleaning. Twelve participants’ data were removed because they did not finish the experiment (i.e., did not complete at least 70% of the daily ESM, or stopped participating altogether).

**Table 1.** Sociodemographic Characteristics of Participants by Condition.

| Characteristic | Emotion Word<br>Training Group | Emotion Word<br>Training Group | Control Word<br>Training Group | Control Word<br>Training Group |
| --- | --- | --- | --- | --- |
|  | *N* =48 | % | *N*=47 | % |
| *Age* |  |  |  |  |
| 18-25 | 23 | 47.92 | 28 | 59.57 |
| 26-31 | 15 | 31.25 | 10 | 21.28 |
| 32-37 | 6 | 12.5 | 0 | 0 |
| 38-43 | 1 | 2.08 | 4 | 8.51 |
| 44-49 | 1 | 2.08 | 2 | 4.26 |
| 50-57 | 0 | 0 | 0 | 0 |
| 58+ | 2 | 4.17 | 3 | 6.38 |
| *Race* |  |  |  |  |
| White | 35 | 67.31 | 36 | 70.59 |
| Hispanic | 1 | 1.92 | 4 | 7.84 |
| Black | 7 | 13.46 | 3 | 5.88 |
| Asian or Pacific Islander | 8 | 15.38 | 8 | 15.69 |
| Native American or Alaska Native | 1 | 1.92 | 0 | 0 |
| *Highest Level of Education Completed* |  |  |  |  |
| High School/GED | 1 | 2.08 | 4 | 8.51 |
| Associate's Degree | 1 | 2.08 | 0 | 0 |
| Bachelor's Degree | 30 | 62.5 | 33 | 70.21 |
| Master's Degree | 16 | 33.33 | 7 | 14.89 |
| Doctoral Degree | 0 | 0 | 3 | 6.38 |
| *Sex Assigned at Birth* |  |  |  |  |
| Female | 36 | 75.0 | 31 | 65.96 |
| Male | 12 | 25.0 | 16 | 34.04 |

*Note.* Descriptive statistics represent all participants. Race is greater than the total number of participants because some participants indicated more than one race.

### Measures

#### Emotion and Control Word Selection

We initially compiled a list of 40 emotions from several sources: the ANEW list [64], an emotion word development list [65], and a .pdf of emotion terms [66]. We then reduced this to a smaller set based on similarity of length and frequency of use in the English language (using [67]). We also selected several emotion words from each quadrant of the affective circumplex [68]. We then piloted the reduced set for understanding, usage, and, in some cases, accuracy in up to three separate pilot samples with differing demographics. Based on the pilot ratings, we then selected two emotion words for each of the seven basic emotions (for the list of ratings, [50, study 5]).

We used a similar process to select control words: We first selected approximately 40 words that were similarly matched for length and frequency of use in the English language to the emotion words. From that list, we piloted the understanding, usage, and (sometimes) accuracy of these words to arrive at a list of 14, which represented normed ratings similar to the chosen emotion words. The overall valence and arousal scores across emotion words were approximately equal to those across the control words.

#### Assessment of Word Knowledge: Emotion and Control Word Understanding, Usage, and Accuracy

We operationalized participants’ knowledge about emotion words and control words through self-report surveys of their usage and understanding of each word and by asking participants to select the correct definition of each word from among four options (see [50]). To measure usage, participants saw the question: “how often do you use this word?” and chose an answer using a 4-point Likert scale: 1 = Often / I typically use this word, 2 = Somewhat / I use this word once in a while, 3 = Rarely / I have used this word, but not regularly, 4 = Never / I don’t think I have ever used this word. To measure understanding, participants saw the question: “how much do you understand this word?”, indicating their answer using a 4-point Likert scale: 1 = Clearly understood / I am fairly confident that I know what this word means, 2 = Possibly understood / I think I know what this word means, but I might be wrong, 3 = Not understood / I don’t know what this word means, but I have heard of it, 4 = I have never heard of this word before. To measure accuracy, participants were asked to select the single correct definition from a list of four definitions.

#### Assessments of Emotional Granularity

We assessed EG using behavioral measures (collected during word training using ESM) and self-reported surveys. For the ESM EG calculation, the average endorsement over the 14-day training period for each of 17 positive and negative feelings was computed for each participant. The feelings were emotion terms (angry, afraid, disgusted, embarrassed, lonely, sad, stressed, tired, worried, calm, happy, excited, grateful, proud, relieved, satisfied, surprised) taken from previous ESM studies (e.g., [37]). Eight feelings were positive and nine were negative. For this study, “surprise” was considered a positive emotion. Using SPSS 29.0, we conducted intraclass correlations (ICCs) on each participant’s average endorsement of the positive feelings and negative feelings during the sampling period. Each participant could have up to 70 occurrences of sampling if they answered each ESM prompt every day of the training. To calculate the ICC for negative and positive emotions for each participant, we used a two-way mixed model reliability analysis with random effects for participant and fixed effects for endorsement measures [32, 45]. The ICC represents the degree to which emotion terms are used interchangeably across instances. By convention, the scores were z-transformed and multiplied by −1 so that higher scores represent greater EG.

Additionally, we used three validated, surveys to measure self-reported EG at the beginning and the end of the experiment. The first survey was the Toronto Alexithymia Scale (TAS) [48]. The TAS 20 contains 20 items that relate to three subscales used to measure externally-oriented thinking and difficulties identifying and describing emotions. Due to an error in our coding, we used different anchors than in the original article (1 = strongly disagree and 5 = strongly agree). Therefore, we removed this entire scale from analysis.

The second survey was the Range and Differentiation of Emotional Experience Scale (RDEES) [47]. The RDEES contains 14 items that relate to two subscales used to measure participants’ range (e.g., “I tend to experience a broad range of different feelings”) and differentiation of emotions (e.g., “I am aware that each emotion has a completely different meaning”). The model has a good fit (RMSEA = 0.06) and reliability (Cronbach’s α = 0.85) [47]. Responses are measured on a 5-point Likert scale: (1) strongly disagree to (5) strongly agree. The RDEES is scored by taking the sum of all items. Four items are reverse-coded. We treated the two subscales as separate mediators.

The third survey was the Trait Meta-Mood scale (TMMS) [54]. The TMMS contains 48 items that relate to three subscales used to measure participants’ mood repair (e.g., “When I become upset, I remind myself of all the pleasures in life”), attention to feelings (e.g., “I think about my mood constantly”), and clarity of feelings (e.g., “I am rarely confused about how I feel”). Each subscale shows good internal consistency, with Cronbach’s α ranging from 0.82 to 0.87 [54]. Responses are measured on a 5-point Likert scale: (1) strongly disagree to (5) strongly agree. Fifteen items are reverse-coded. We treated all three subscales as separate mediators.

#### Assessments of Emotional Regulation, Dysregulation, and General Well-Being (Mental Health, and Satisfaction with Life)

We used the following four validated, self-reported surveys for each of the three outcome variables.

For emotion regulation, we used the Emotional Regulation Questionnaire; ERQ) [59]. The ERQ contains 10 items that relate to two subscales assessing participants’ emotional reappraisal (e.g., “I control my emotions by changing the way I think about the situation I’m in”) and emotional suppression (e.g., “I keep my emotions to myself”). The ERQ shows acceptable to excellent reliability with a Cronbach’s α between 0.76 and 0.90, and strong concurrent validity. Responses are measured on a 7-point Likert Scale: (1) strongly disagree to (7) strongly agree. The ERQ is scored by taking the average across all items from both scales. We treated the subscales as separate outcomes.

To assess emotional dysregulation, we used the Difficulties in Emotion Regulation (DERS) scale [60]. The DERS contains 18 items that measure six areas of emotional dysregulation (i.e., clarity, awareness, goal-orientation during distress, strategies, ability to control impulses, and non-acceptance of emotions). An example of a question is: “When I’m upset, I feel guilty for feeling that way”. The DERS has previously established excellent internal consistency (Cronbach’s α = 0.92), test-retest reliability, and both divergent and discriminant validity [69]. Responses are measured on a 5-point scale: (1) almost never (5) almost always. The DERS is scored by taking the sum of all items. Three items are reverse-coded. We used the entire scale as an outcome.

To measure well-being, we used two surveys. First, we used the Mental Health Continuum – Short Form; MHC-SF) [62]. The MHC-SF contains 14 items adapted from the 40-item Mental Health Continuum – Long Form that assesses participants’ well-being across three domains (i.e., psychological, emotional, and social). Responses are measured on a 6-point Likert scale: (1) never to (6) every day. An example question is: “In the past month, how often did you feel happy?” The MHC-SF has demonstrated excellent reliability as a whole (Cronbach’s α = 0.89), and acceptable convergent (correlations between 0.17 and 0.49) and discriminant validity (0.07 and 0.39) [62]. For this study, we used the sum across the three domains for a total picture of “well-being”.

Lastly, we also used the Satisfaction with Life Scale (SWL) as our second measure of well-being [61]. The SWL is a 5-item measure that measures responses on a 7-point Likert scale ranging from (1) strongly disagree to (7) strongly agree. The SWL assesses participants’ well-being and satisfaction with life as a whole (e.g., “The conditions of my life are excellent”) [61]. The SWL is reliable, with a Cronbach’s α of 0.87, and has excellent test-retest reliability and discriminant validity [70]. This scale is scored by taking the sum of all items.

#### Training Materials

The approach for this study was to train two groups to either increase their emotion word knowledge or control word knowledge. The emotion word group received training materials for 14 highly-precise emotion words, whereas the control word group received training materials for 14 length- and frequency-matched control words. The 14 emotion words and 14 control words were the same as those presented in the baseline word knowledge assessments. The training materials consisted of 1) a definition of the word, 2) a scenario describing the meaning of the word, 3) a cartoon depicting the meaning of the word, 4) a multiple-choice question in which participants had to choose the one sentence that correctly used the word among five sentences, and 5) a disambiguation paragraph explaining how the word was different from related words (e.g., horrified vs. terrified vs. fearful) (Appendix A; B). We chose definitions for the emotion and control words based on those from Merriam-Webster online dictionary, Urban Dictionary, and dictionary.com. We altered definitions if they included another emotion word or if the definition was too similar to the definition of a related word. We created the word scenarios based on those in [52]. We adapted cartoons from emotiontypology.com, a website that provides detailed information on positive and negative emotion words, with permission from the authors. For the multiple-choice question, we created incorrect sentences by substituting the training word into the sentences that would be accurate for other similar words. The multiple-choice question was used to assess engagement with the training materials (see Procedure), and we provided feedback on the correctness of the selection. All final materials and modifications were based on the consensus of the researchers in the lab.

### Procedure

#### General Procedure

The experiment was divided into five stages, spanning 22-26 days, based on the scheduling of Session 2 (see below). We assigned participants to one of the word learning groups using every-other allocation.

#### Session 1

Participants reported in person to the first author’s laboratory. After signing a written consent form, participants were assigned a subject number for confidentiality, which also linked their ESM data to their survey data.

Session 1 consisted of two emotion perception tasks (not reported in this paper) and a demographic questionnaire. All participants then completed a computer survey that showed each of the 14 highly precise emotion words in a random, but pre-determined, order. For each word, participants first indicated their usage and understanding. They were then asked to select the correct definition of the word among four choices. Participants were only asked to indicate their understanding, and subsequently the definition, if they indicated a “1” or “2” for the usage question. If they indicated a “3” or “4”, meaning that they did not use or only rarely used the word, we did not ask about their understanding or whether they could select the accurate definition, as it would likely not yield reliable information. After rating all 14 emotion words, participants repeated the same procedure for the 14 control words (also in a random but predetermined order).

Participants then completed the following surveys in this order: RDEES, TAS-20, DERS, TMMS, ERQ, MHC-SF, and SWL. After completing the surveys, participants downloaded M-Path (for ESM sampling; www.m-Path.io) on their smartphone and completed a short sample of the ESM procedure. The total time to complete Session 1 took between 45 and 60 minutes.

#### Pre - ESM Recording Period, Training Period, and Post–ESM Recording Period

The first four days following Session 1 constituted the pre-ESM sampling period. Participants received five “samples” per day at 9 am, 12 pm, 3 pm, 6 pm, and 9 pm, Central Standard Time. Participants had up to one hour to respond to the sample. Participants saw 17 basic feeling words (e.g., happy, sad, etc.) in a predetermined order and were asked to rate how intensely they were currently feeling each using a 5-point Likert scale (1 = Extremely to 5 = Not at All). None of the presented feeling words were the highly precise emotion words used in the training materials. Each sampling took about two minutes to complete, for a total of about 10 minutes per day.

The 14-day training period began the following day. One hour before the first ESM sampling, participants received a notification through the M-path app containing a training word with its training materials. At the end of the materials, participants were asked to answer the multiple-choice question by selecting the correct usage of the word in a sentence from among five options. Participants received feedback on their responses and explanation of how that word differed from related words. The delivery, reading, and response to this question added about 10 minutes each day.

The four days following the training period were the post-ESM sampling period. Participants completed the same five samples they had completed in the pre-ESM period. We did not analyze ESM data during the pre- or post-period. The pre-period was to ensure that participants were familiar with the sampling before introducing the training words, and the post-period was used to ensure that participants continued to engage in the experiment until they returned for Session 2.

### Session 2

Participants completed the same measures as in Session 1 in the laboratory. Based on scheduling availability, participants completed Session 2 anytime within the post-sampling period up to two days after. Participants received a hard copy of all training word materials for their use during their debriefing, regardless of which condition they were assigned. The total time to complete Session 2 was between 30 and 40 minutes.

## Data Processing

To test our hypotheses, we performed mediation analyses separately for the emotion word training group (n = 46) and the control word learning group (n = 46), since we only expected that EG measures would mediate the relationship between word knowledge variables and outcome measures in the emotion word group. We had no a priori prediction about whether there would be a mediation or difference in outcomes for the control word group. For **each** group, we performed <u>two</u> robust, mediated analyses: one used the self-reported EG measures as mediators, and the other used the behaviorally-derived EG estimates from the ICC calculations as mediators. All analyses were conducted in JASP (v. 19.01), using standardized estimates for all variables, with auto estimate and no emulation and 1000 replication bootstrapping (bias-corrected). For the mediated analyses for the emotion word learning group, we used control word knowledge variables as background confounders; whereas for the control word learning group, we used the emotion word knowledge variables as background confounders.

### Data Analysis Plan and Reduction of Variables

All variables in the mediated analysis were based on the difference in performance between baseline (pre, Session 1) and completion (post, Session 2), with the exception of ICCs, which were calculated over the training period. From our candidate list of variables to include in the model, we performed variable selection for inputs, mediators, and outcomes by assessing correlation structures and conducting EFAs, eliminating variables that were highly correlated or loaded onto the same factors. The TAS scores were not reported due to an error in the anchors of the scale, and emotion perception variables are reported in another manuscript. All other manipulations and data are reported in the following (see also Supplemental Analyses).

### Factor Structure and Correlations of Variables for Models

#### Input Variables

We found word accuracy and usage to be negatively correlated for both emotion (*r* = - 0.323, *n = 96, p* < 0.001, 95% C.I. [−0.492, −0.131], Fischer’s z = −0.335) and for control words (*r =* - 0.581, *n = 96, p* < 0.001, 95% C.I. [−0.700, 0.430], Fischer’s z = −0.663). We found additional significant correlations among control word and emotion word variables (Supplemental Materials). We used an exploratory factor analysis (EFA; oblique Promax rotation) and found a two-factor structure. Control and emotion word accuracy, and control and emotion word usage meaningfully loaded onto one factor (Eigenvalue = 2.250, rotated squared sum loading = 1.757, proportion variance = 0.293, cumulative variance = 0.293), whereas control and emotion word understanding loaded onto a second factor (Eigenvalue = 1.444, rotated squared sum loading = 1.030, proportion variance = 0.172, cumulative variance = 0.465). Based on these correlations and the observed factor structure, we kept word usage and word understanding in the model. Thus, emotion and control word accuracy were removed as predictor variables in the mediated models.

#### Mediator Variables

Because the TAS was incorrectly anchored, we removed the TAS from the analyses and only analyzed the TMMS subscales and the RDEES subscales. We found a significant correlation between the RDEES Range (RDEES_R) and Differentiation (RDEES_D) subscales (*r =* 0.241, n = 96, *p* = 0.018, 95% C.I. [0.043, 0.421], Fischer’s z = 0.246). We found an additional significant correlation between the TMMS Clarity of Feelings (TMMS_CF) and Mood Repair (TMMS_MR) (*r =* 0.245, n = 96, *p* = 0.016, 95% C.I. [0.047, 0.424], Fischer’s z = 0.250). We found a two-factor structure using an EFA: one containing TMMS_MR (Eigenvalue = 1.400, rotated sum square loading = 0.687, proportion variance = 0.137), and a second containing RDEES_R and RDEES_D (Eigenvalue = 1.221, rotated sum square loading = 0.579, proportion variance = 0.116). The TMMS_CF and TMMS_AF did not load onto a factor with an eigenvalue greater than 1.0. We elected to keep only TMMS_MR and the RDEES_D as the self-reported EG measures in our model. Therefore, we removed two subscales of the TMMS and one of the RDEES as mediator variables in our model.

We found that ICCs for the negative and positive emotion words were statistically correlated (*r* = 0.388, *n = 96, p <* 0.001, C.I. [0.203, 0.546], Fisher’s z = 0.406) and loaded onto one factor using an EFA (Eigenvalue = 1.388, rotated sum square = 0.775, proportion variance = 0.388). We chose to keep the ICC for negative words as the behaviorally-based measure of EG. The ICCs from the negative emotions were not significantly correlated with any self-reported EG scales. Therefore, we did not use the ICCs of the positive emotion words in our model.

#### Output Variables

We found that the SWL outcome was significantly correlated with the MHC-SF (*r* = 0.440, n = 96, *p* < 0.001, 95% C.I. [0.263, 0.589], Fischer’s z = 0.472) and the DERS (*r =* - 0.215, n = 96, *p* = 0.036, 95% C.I. [−0.398, −0.015], Fischer’s z = −0.218). Using an EFA, we found two factors: SWL and the MHC-SF loaded onto one factor (Eigenvalue = 1.602, rotated sum square loading = 1.038, proportion variance = 0.208), and the ERQ_S loaded onto a second factor (Eigenvalue = 1.033, rotated sum square = 0.999, proportion variance = 0.200). The DERS and the ERQ_R did not load onto either factor. Yet, since the DERS was only mildly correlated with the SWL, we kept it as an output variable alongside the SWL and ERQ_S. Therefore, we did not use the ERQ_R or the MHC as outcome variables in our model.

## Results

### Emotion Word Group (n = 48)

#### Change in Measures Post-Pre

Emotion word usage and understanding improved after training (Table 2), although the results were not statistically significant using traditional dependent t-tests. Bayesian analyses, however, showed that the change in results from both variables was within 6% error for emotion usage (95% C.I. [1.95-2.20 pre-training] [1.89-2.19 post-training]; logBF10 = −1.49) and for emotion understanding (95% C.I. [1.11-1.22 pre-training] [1.08-1.21 post-training]; logBF10 = - 1.64). Participants in the emotion word group also showed improved differentiation of emotional experiences (RDEES _D), decreased emotional dysregulation (DERS), decreased suppression (ERQ_S), increased mood repair (TMMS_MR), and a slight improvement in the satisfaction with life (SWL) after training, although again not statistically significant with traditional statistics. The Bayes change value for the DERS was within 5% error (95% C.I. [35.03-40.85 pre-training] and [33.86-39.10 post-training]; logBF_10_ = −0.995). Bayes values were also within 6% error for the RDEES_D (95% C.I. [24.12-27.13 pre-training] [24.59 – 27.36 post-training]; logBF_10_ = −1.578) and for the TMMS_MR (95% C.I. [38.19-40.48 pre-training] [38.62-41.01 post-training]; logBF_10_= −1.489). The Bayes value for SWL was within 7% error (95% C.I. [21.49-25.47 pre-training] [21.39-25.65 post-training]; logBF_10_ = −1.845). Finally, the Bayes value for ERQ_S was within 4% error (95% C.I. [2.97-3.69 pre-training] [2.81-3.5 post-training]; logBF_10_ = −0.796). Surprisingly, learning emotion words also resulted in *improved use and understanding of control words,* which we did not predict, though this change was not statistically significant with traditional statistics.

**Table 2.**
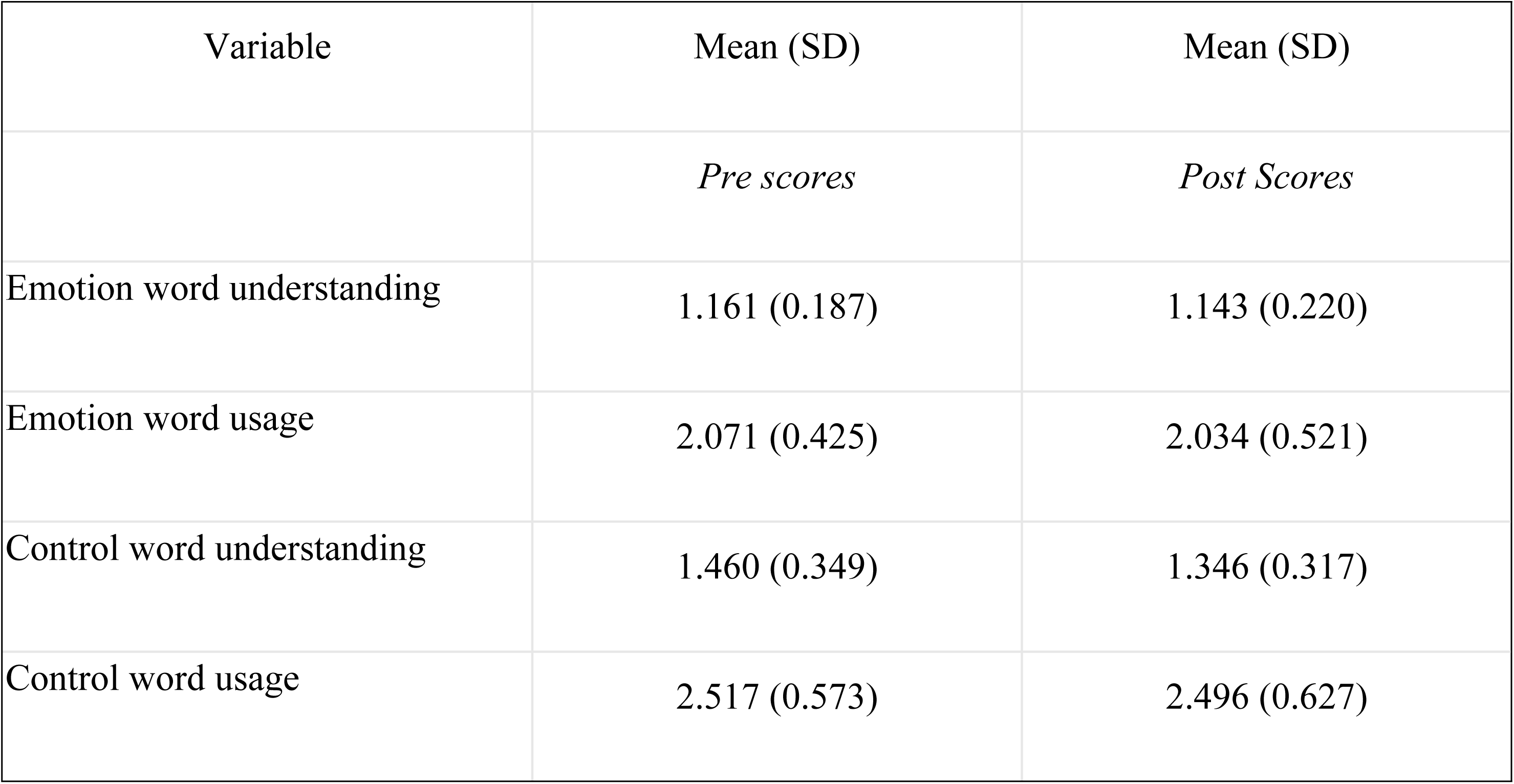
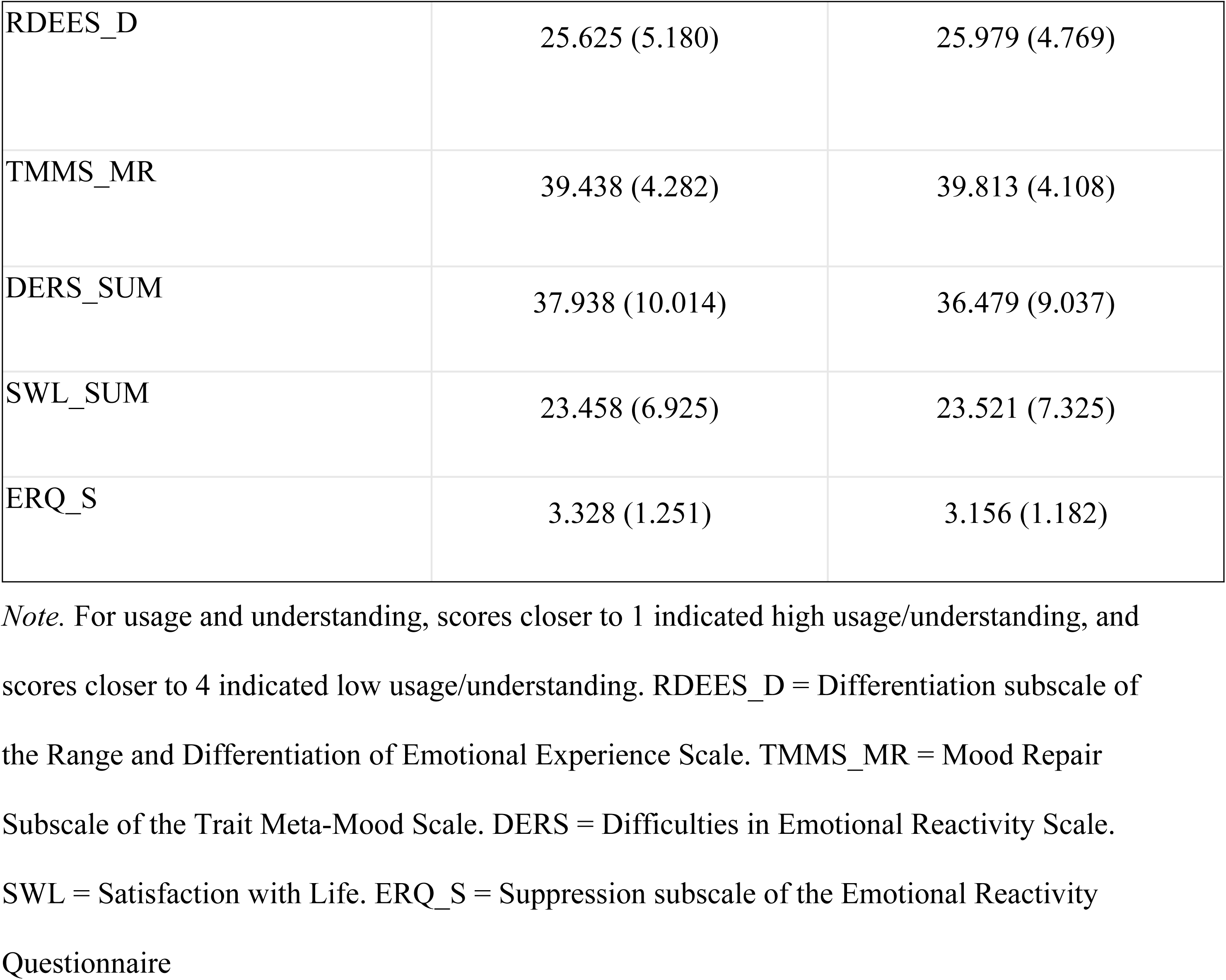
Descriptive Statistics for Measures Pre and Post for Emotion Word group.

| Variable | Mean (SD) | Mean (SD) |
| --- | --- | --- |
|  | *Pre scores* | *Post Scores* |
| Emotion word understanding | 1.161 (0.187) | 1.143 (0.220) |
| Emotion word usage | 2.071 (0.425) | 2.034 (0.521) |
| Control word understanding | 1.460 (0.349) | 1.346 (0.317) |
| Control word usage | 2.517 (0.573) | 2.496 (0.627) |
| RDEES_D | 25.625 (5.180) | 25.979 (4.769) |
| TMMS_MR | 39.438 (4.282) | 39.813 (4.108) |
| DERS_SUM | 37.938 (10.014) | 36.479 (9.037) |
| SWL_SUM | 23.458 (6.925) | 23.521 (7.325) |
| ERQ_S | 3.328 (1.251) | 3.156 (1.182) |

*Note.* For usage and understanding, scores closer to 1 indicated high usage/understanding, and scores closer to 4 indicated low usage/understanding. RDEES\_D = Differentiation subscale of the Range and Differentiation of Emotional Experience Scale. TMMS\_MR = Mood Repair Subscale of the Trait Meta-Mood Scale. DERS = Difficulties in Emotional Reactivity Scale. SWL = Satisfaction with Life. ERQ\_S = Suppression subscale of the Emotional Reactivity Questionnaire

#### Mediated Model Results

The first model used the reduced set of self-reported measures of EG as mediators. There were no significant direct or indirect effects. There was one marginally significant total effect between emotion word usage and the DERS (estimate = −0.312, SE = 0.162, z-value = −1.921, *p* = 0.055, 95% C.I. [−0.687, −0.011]). A few other paths were significant (Figure 1): 1) from the RDEES_D to the DERS (estimate = −0.426, SE = 0.125, z-value = −3.398, *p* < 0.001, 95% C.I. [−0.687, −0.199]) and 2) from the RDEES_D to the ERQ_S (estimate = −0.278, SE = 0.135, z-value = −2.066, *p* = 0.039, 95% C.I. [−0.628, −0.018]). Additionally, the pathway from emotion word usage to the RDEES_D was significant (estimate = 0.341, SE = 0.168, z-value = 2.030, *p* = .042, bias-corrected 95% C.I. [−0.033, 0.691]). Significant pathway effects for the background confounders are not reported.

**Figure 1.**
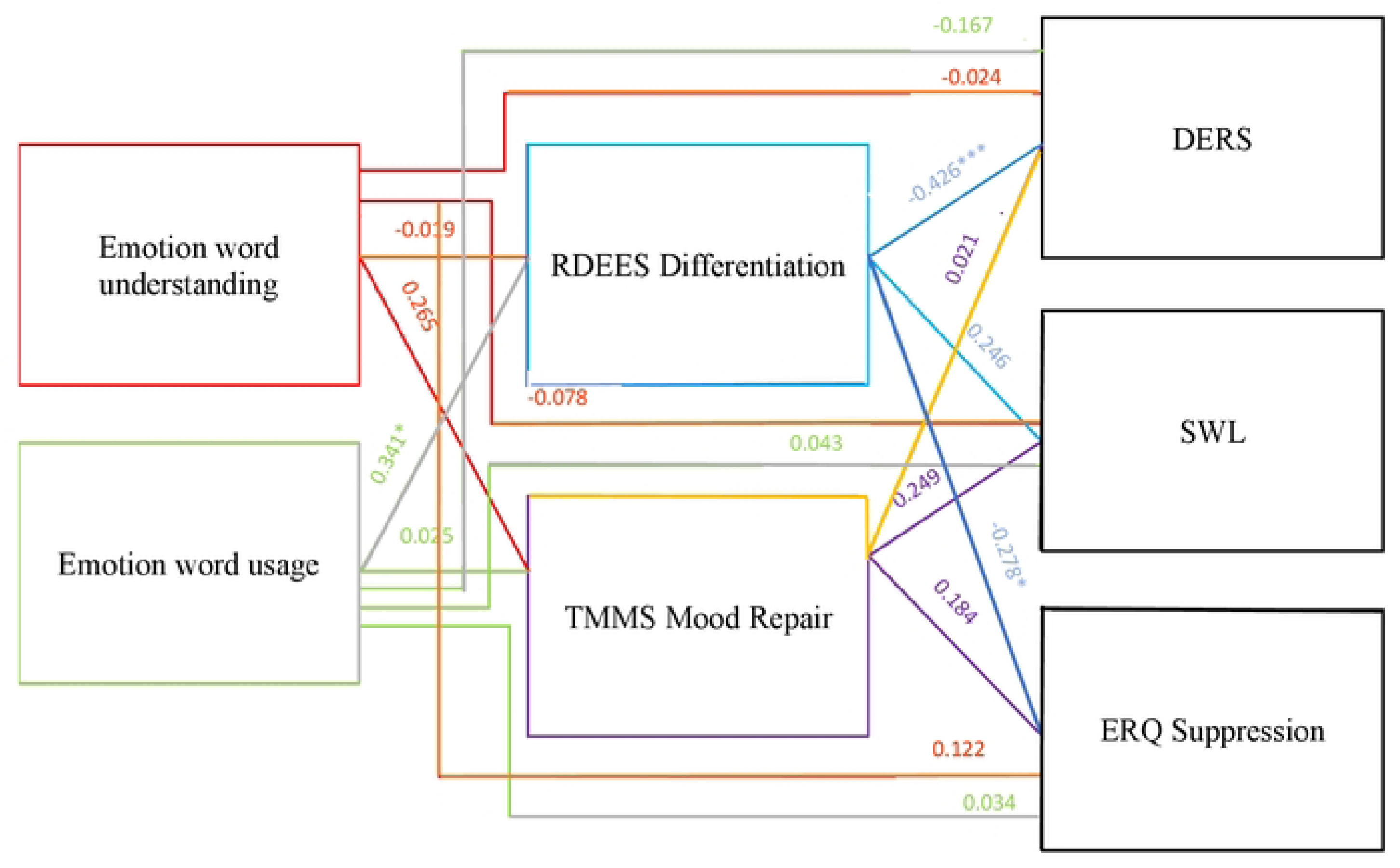
Mediated Model of Emotion Work Group with Self-Reported EG Measures as Mediators. ***Note.*** Control pathways are not shown. Only path coefficients between variables are shown. Coefficients are z-values. * p<.05, *** p<.001.

The second model used the behavioral measure of EG (derived from the ICCs) as the mediator. There were no significant indirect or total effects, although there was one marginally significant direct effect between emotion word usage and the DERS (estimate = −0.313, SE = 0.162, z-value = −1.928, *p* = 0.054, 95% C.I. [−0.683, −0.017]) (Figure 2). Thus, unlike the self-reported EG mediators, there were no similar mediating effects when using the ICCs from negative emotion words. Significant pathway effects for the background confounders are not reported.

**Figure 2.**
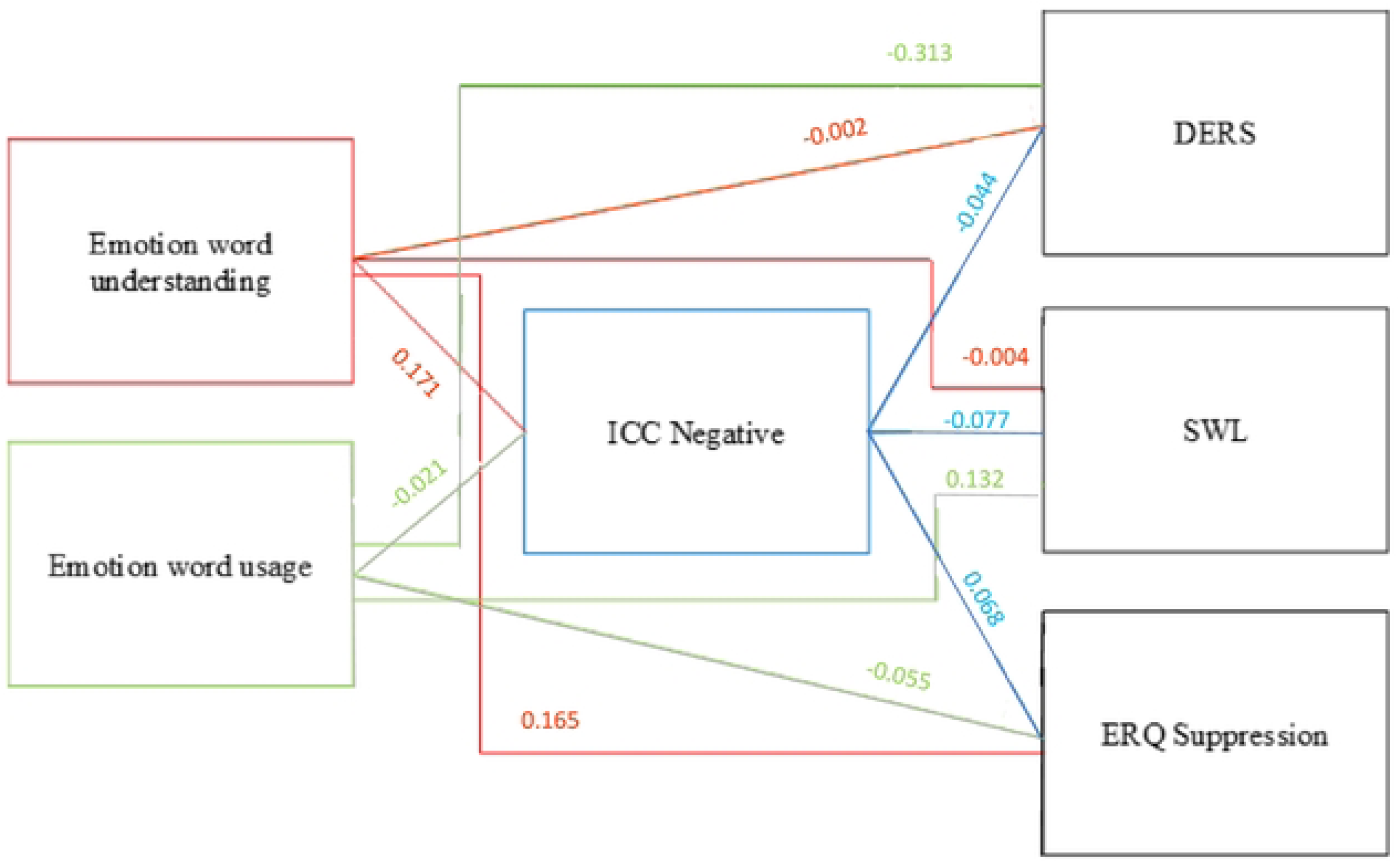
Mediated Model of Emotion Word Group with ICCs as Mediator. ***Note.*** Control pathways are not shown. Only path coefficients between variables are shown. Coefficients are z-values. No pathways are significant at any level.

### Control Word Group (n = 47)

#### Change in Measures Post-Pre

As expected, control word usage and understanding improved after training for the control group (Table 3), but only significantly so for control word usage, t(47) = 1.847, *p* = 0.035, Cohen’s d = 0.267, with Bayes analysis under 3% error (95% C.I. [2.43-2.74 pre-training] [2.32-2.60 post-training]; logBF_10_ = −0.286). The Bayes error for control word understanding was under 6% (95% C.I. [1.389-1.588 pre-training] [1.34-1.56 post-training]; logBF_10_ = −1.684). Interestingly, the differentiation of emotional experiences (RDEES_D) *decreased* after training (95% C.I. [24.61-26.98 pre-training] [24.29-26.70 post-training]; logBF_10_ = −1.789, error under 7%), whereas emotional dysregulation (DERS) *increased* after training (95% C.I. [31.62-36.26 pre-training] [32.04-36.79 post-training]; logBF_10_ = −1.647, error under 6%). These results clearly differed from what was observed in the emotion word training group. However, similar to what was observed in the emotion word training group, mood repair (TMMS_MR) and satisfaction with life (SWL) improved after training (95% C.I. [39.37-41.76 pre-training] [40.42-42.21 post-training]; logBF_10_ = −0.899, error at 4%); (95% C.I. [25.89-28.86 pre-training] [26.36-29.18 post-training]; logBF_10_ = −1.547, error under 6%), and suppression (ERQ_S) decreased after training (95% C.I. [3.00-3.66 pre-training] [2.91-3.55 post-training]; logBF10 = −1.430, error 6%). Interestingly, learning control words also resulted in *increased emotion word use,* but *worse understanding of emotion words*, neither of which was predicted.

**Table 3.** Descriptive Statistics for Measures Pre and Post for Control Word group.

| Variable | Mean (SD) | Mean (SD) |
| --- | --- | --- |
|  | *Pre scores* | *Post Scores* |
| Control word understanding | 1.488 (0.342) | 1.449 (0.376) |
| Control word usage | 2.585 (0.532) | 2.458 (0.492) |
| Emotion word usage | 2.066 (0.405) | 1.929 (0.367) |
| Emotion word understanding | 1.203 (0.257) | 1.253 (0.274) |
| RDEES_D | 25.792 (4.079) | 25.646 (4.656) |
| TMMS_MR | 40.563 (4.115) | 41.313 (3.075) |
| SWL_SUM | 27.375 (5.110) | 27.771 (4.865) |
| ERQ_S | 3.328 (1.128) | 3.229 (1.117) |
| DERS_SUM | 33.938 (7.996) | 34.417 (8.181) |

*Note.* For usage and understanding, scores closer to 1 indicated high usage, and scores closer to 4 indicated low usage. RDEES\_D = Differentiation subscale of the Range and Differentiation of Emotional Experience Scale. TMMS\_MR = Mood Repair Subscale of the Trait Meta-Mood Scale. DERS = Difficulties in Emotional Reactivity Scale. SWL = Satisfaction with Life. ERQ\_S = Suppression subscale of the Emotional Reactivity Questionnaire

#### Mediated Model Results

When using the reduced set of self-reported measures of EG, we observed a significant direct effect between control word usage and SWL (estimate = 0.374, SE = 0.152, z-value = 2.466, *p* = 0.014, 95% C.I. [0.034, 0.804]) alongside a significant total effect (estimate = 0.406, SE = 0.156, z-value = 2.599, *p =* 0.009, 95% C.I. [0.086, 0.805]). There were no significant indirect effects, and no other pathways were significant (Figure 3).

**Figure 3.**
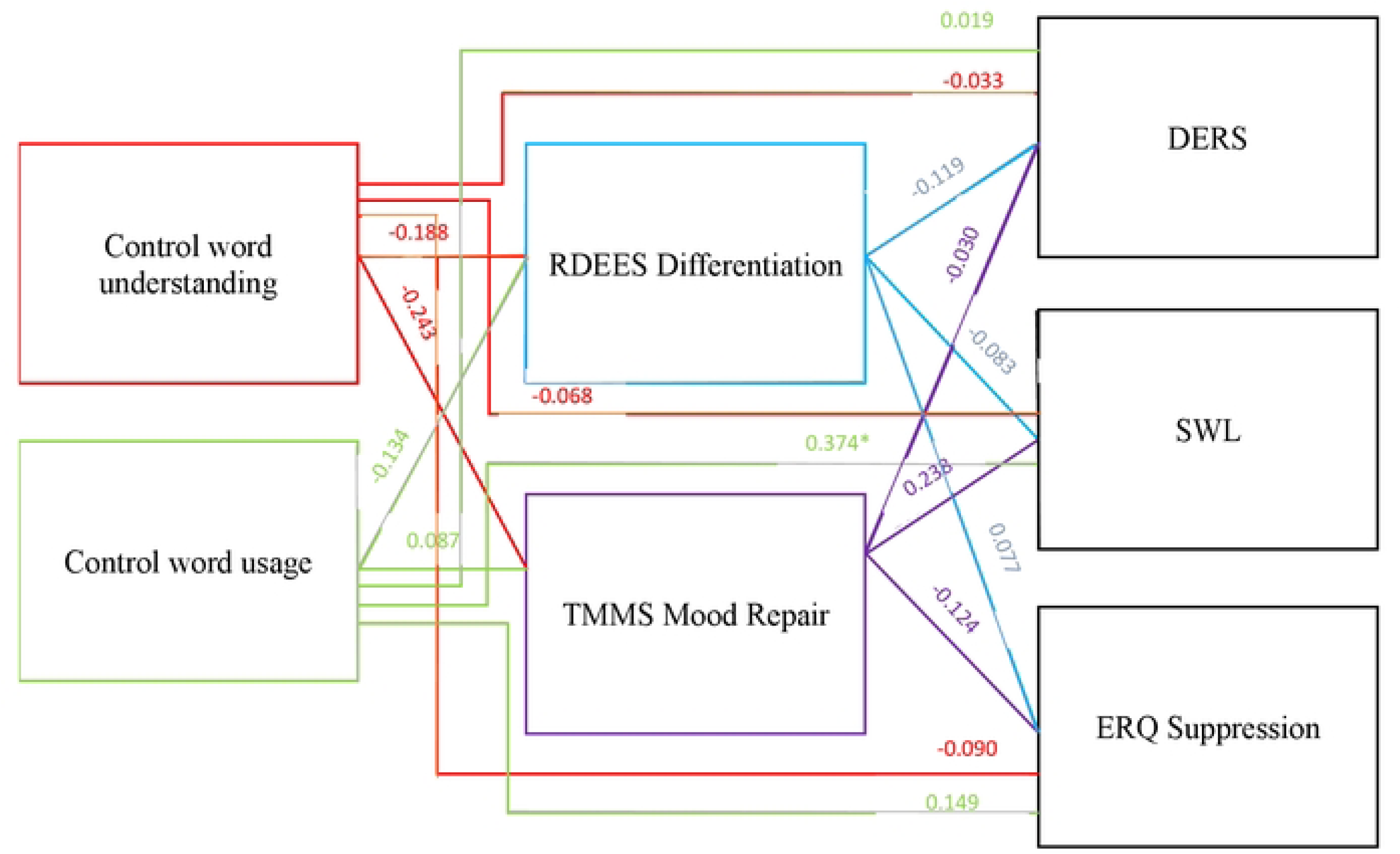
Mediated Model of Control Word Group with Self-Reported EG Measures as Mediators. ***Note.*** Control pathways are not shown. Only path coefficients between variables are shown. Coefficients are z-values.* *p*<.05.

In the second mediated analysis, which used the behavioral measure of EG (derived from the ICCs) as a mediator, there was one significant direct effect between control word usage and the SWL, similar to that above (estimate = 0.406, SE = 0.154, z-value = 2.640, *p =* 0.008, 95% C.I. [0.111, 0.792]). In addition, there was also a significant total effect between the two (estimate = 0.407, SE = 0.156, z-value = 2.603, *p* = 0.009, 95% C.I. [0.114, 0.833]). There was no significant indirect effect, and no other additional pathways were significant (Figure 4).

**Figure 4.**
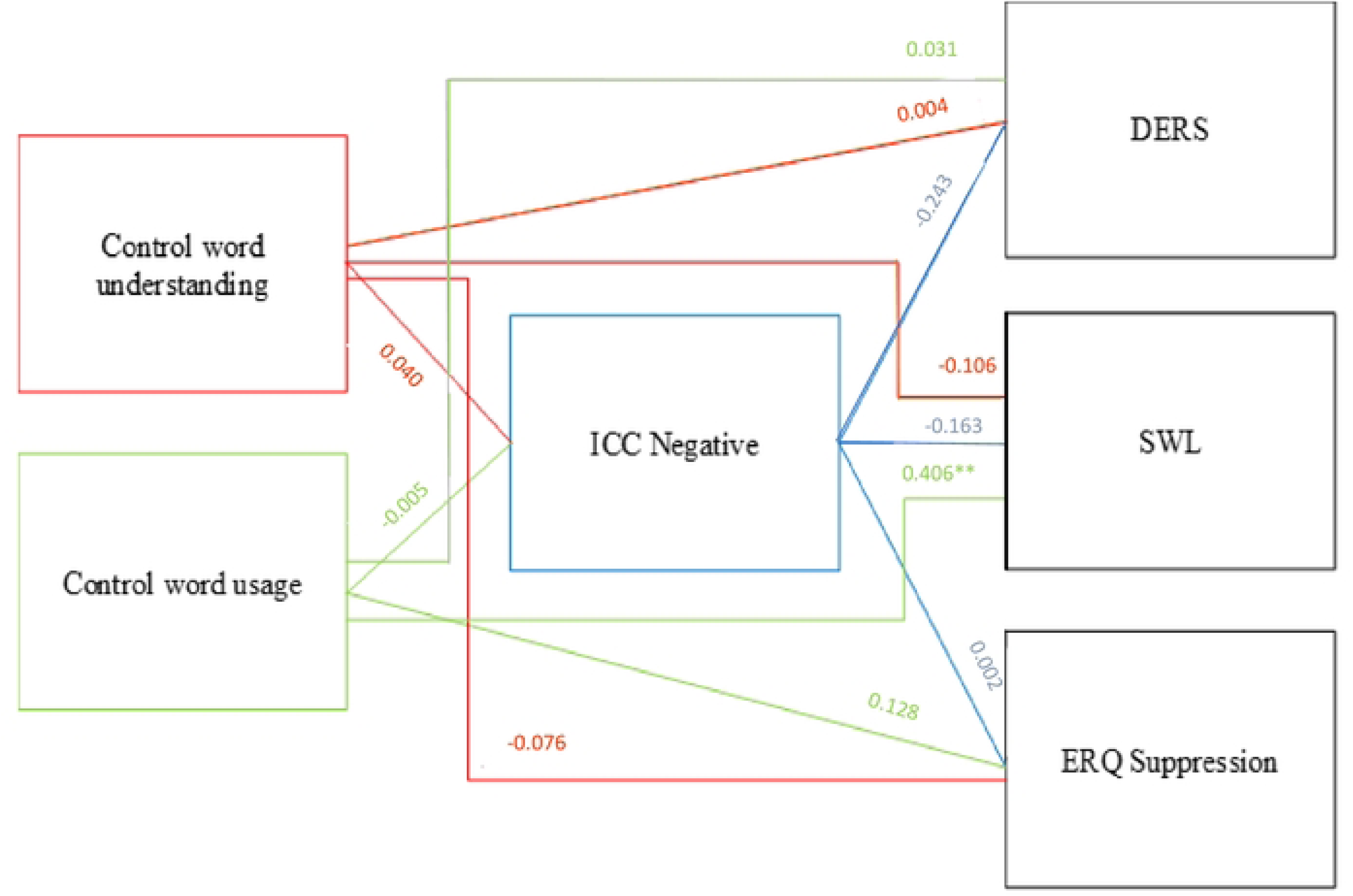
Mediated Model of Control Word Group with ICCs as Mediator. ***Note.*** Control pathways are not shown. Only path coefficients between variables are shown. Coefficients are z-values. * *p*<.05, ** *p*<.001.

### Exploratory Modeling: Lasso Models

Because the planned mediation models yielded limited evidence for any EG measures as mediators, we conducted a series of exploratory Lasso regression analyses to understand any additional relationships that may exist between variables. Lasso regression is a regularization technique that performs variable selection by penalizing model complexity through shrinking variable parameters to zero. This dimension reduction prevents overfitting and addresses issues of collinearity. The result is a method that is well-suited for exploratory analyses when presented with correlated predictors [72]. Thus, the variables that were excluded from the previous mediated models after considering correlations and factor analysis were included in Lasso regressions.

In our full collection of data, all individuals had measures from six word-related variables: the change in their self-reported ability to (1) understand, (2) use, and (3) accurately pick the correct definition for emotion words, and to (4) understand, (5) use, and (6) accurately pick the correct definition for control words. Individuals also had measures from six EG-related variables: (1) intra-class correlations among negative emotion words from the ESM (ICC_Neg and ICC_Pos), (2) range (RDEES_R) and (3) differentiation (RDEES_D) of emotional experiences, and (4) mood repair (TMMS_MR), (5) clarity of feelings (TMMS_CF), and (6) attention to feeling (TMMS_AF) of trait meta-mood. Participants also had five outcome measures: (1) emotional regulation suppression (ERQ_S) and (2) emotional regulation reappraisal (ERQ_R), (3) emotional dysregulation (DERS), (4) satisfaction of life (SWL), and (5) mental health (MHC-SF). Finally, participants were also assigned to the emotion word or control word training group, which served as a grouping variable.

As these models are exploratory, we considered multiple models for each of the five outcomes. We first performed Lasso regression using one EG variable at a time, alongside all word-related variables and the grouping variable (treatment group), allowing no interactions among the independent variables. Six models (one for each EG variable) were fit per outcome in this scenario. We then performed the regressions allowing interactions between the EG variable and the independent variables; this scenario also produced six models per outcome. Next, we performed Lasso regression on a full model containing all six EG measures, allowing all to interact with any independent variable. There was one model per outcome in this scenario. Finally, we performed Lasso regression with two additional scenarios per outcome. Considering the self-reported measures as sets, we fit (1) one model allowing the three TMMS measures to interact with the independent variables and (2) another allowing the two RDEES measures to interact with the independent variables.

In all, we performed Lasso regression on 15 models for each outcome. Below, we report the models that explained greater than 10% of the variance in the outcome (*R*^2^ ≥ .10). When interpreting models, it is important to remember that these regressions do not imply causal structure and no pathways are being tested. Therefore, the variables do not have the conceptual assignments of “predictors” and “mediators”. All Lasso models are reported in the Supplementary Analyses after the JASP-mediated models.

#### Lasso Model on DERS outcome

Five of the fifteen models explained at least 10% of the observed variance in emotional dysregulation (DERS). The model explaining the most variation (R^2^ = 0.335) contained all six EG measures and allowed each to interact with all independent variables. The other four models had R^2^ values ranging from 0.103 (the model with no interactions) to 0.326 (allowing interactions from the set of three TMMS measures). In the best model, the grouping variable (emotion or control word group) was removed during regularization, indicating group differences did not meaningfully change the DERS outcome after accounting for the other predictors and their interactions. The variables retained were control word accuracy, RDEES_D, TMMS_CF, and several interactions between these and word knowledge (emotion word understanding with TMMS_AF and TMMS_MR; emotion word usage with TMMS_AF, TMMS_MR, and RDEES_D; emotion word accuracy with RDEES_R and TMMS_MR; control word understanding with ICC_Neg; control word usage with TMMS_AF; control word accuracy with TMMS_CF) (see Supplementary Analyses for the estimated effects). The findings revealed that emotion word usage, understanding, accuracy interacted with various aspects of the TMMS and RDEES, and control word usage and accuracy interacted with only aspects of the TMMS. In addition, control word understanding interacted only with the behaviorally-derived measure of EG (ICC_Neg).

#### Lasso Model on ERQ Reappraisal Outcome

The model explaining most of the variation in ERQ_R allowed the two measures of the RDEES to interact (R^2^ = 0.171) with the word knowledge variables. Two other models also explained at least 10% of the observed variation in ERQ_R: (1) the full model containing all six EG measures and their interactions with word knowledge variables (R^2^ = 0.141), and (2) the model that allowed interaction with the singular RDEES_D measure (R^2^ = 0.162). In all these models, the grouping variable was removed. In the best model, the retained variables were the RDEES_R and an interaction between emotion word usage and the RDEES_D.

#### Lasso Model on ERQ Suppression Outcome

There were no models explaining at least 10% of the observed variance in ERQ_S.

#### Lasso Model on SWL Outcome

The model explaining most of the variation in the SWL (R^2^ = 0.126) allowed the three TMMS variables to interact with the remaining independent variables. The only other model that explained more than 10% of the observed variation allowed interactions only from the TMMS_MR measure (R^2^ = 0.124). Again, in these models, the grouping variable was removed after accounting for the other predictors and their interactions. In the best model, the variables retained were the TMMS_MR, TMMS_CR, and an interaction between TMMS_MR and emotion word and control word usage.

#### Lasso Model on MHC Outcome

The model that explained most of the observed variation in the MCH contained only the TMMS_MR EG variable (R^2^ = 0.112). This model retained word group as a meaningful variable, highlighting potential differences in changes in the mental health outcome between the two groups. Additional variables retained in this model included TMMS_MR, emotion word accuracy and understanding, and control word usage and understanding.

## Discussion

This study examined whether increasing word knowledge (learning either specific emotion words or control words) affected emotion regulation, dysregulation, and overall well-being, while exploring the possible mediating roles of various EG measures. Specifically, we collected both self-reported survey measures of EG and behaviorally-derived measures of EG. Word knowledge was operationalized as changes in participants’ usage, understanding, and ability to accurately define training words in the group to which they were assigned (emotion word or control word). Increases in word knowledge for words in the opposite training group were included in the mediation models as background confounds to highlight changes in only the words that were taught to each group.

### Proof of Concept: Training

Participants trained in <u>emotion words</u> showed improved usage and understanding of emotion words after training (but also increases in the use and understanding of *control words*). Although results did not reach statistical significance, the results show initial positive proof-of-concept for this type of training paradigm.

Participants assigned to emotion words also showed slightly improved emotional differentiation (RDEES_D), increased mood repair (TMMS_MR), increased satisfaction with life (SWL), lower emotional dysregulation (DERS), and less suppression (ERQ_S) after training. Therefore, emotion word training affected all outcomes in the predicted (desirable) direction. Although the changes were small in some cases and did not reach statistical significance, the results show that not only does emotion word training increase knowledge for these words, but also that it produces positive changes in EG with improved emotion regulation and well-being outcomes.

Participants assigned to learn <u>control words</u> showed increased usage and understanding of control words after training, although most effects did not reach traditional levels of statistical significance. They also showed some improvement in *emotion word* usage but decreases in *emotion word* understanding after training. Participants assigned to this condition also showed some increases in mood repair (TMMS_MR), satisfaction with life (SWL), and emotional dysregulation (DERS), but decreases in the use of suppression (ERQ_S) and in emotional differentiation (RDEES_D). Therefore, training using control words had some desirable effects (e.g., decreased suppression and improved satisfaction with life and mood repair) and some undesirable effects (e.g., decreased EG and increased emotional dysregulation).

### Summary of Mediated and Lasso Effects

For those participants assigned to learn <u>emotion words</u>, we found that an increased ability to differentiate emotional experiences on self-reported measures of EG was associated with decreased emotional dysregulation and also decreased suppression, both of which are adaptive outcomes. We also found that as emotion word usage increased, self-reported EG (RDEES_D) increased. We did not find strong evidence that self-reported measures of EG mediated the relationship between emotion word usage and dysregulation or suppression, nor did we see any significant mediated effects or partial pathways when using behaviorally-derived measures of EG. Additionally, the finding that the behaviorally-derived and the self-reported estimates of EG were not highly correlated provides some insight that they do not target and/or do not refer to the same ability, but rather potentially related individual aspects of a larger construct. Perhaps it is people’s *beliefs* (captured from survey measures) that mediate the relationship between emotion word usage and dysregulation rather than their *actual* emotional differentiation (captured from ICC estimates). These results are consistent with our Lasso models that showed emotion word usage, understanding, and accuracy interact with various aspects of self-reported EG, but not with behavioral measures of EG, when considering emotional dysregulation. We also found that emotion word usage interacted with RDEES_D to affect the outcome of emotional reappraisal, as well as with TMMS_MR to affect the outcome of satisfaction with life. It is important to remember, however, that word training group was not retained in any of the aforementioned Lasso analyses, which implies that these relationships hold for individuals in both learning and control word groups. Finally, these models showed that emotion word accuracy and understanding had simple effects on mental health. In the case of mental health, however, word training group was retained, suggesting that the effects of emotion word accuracy and understanding affected mental health differently based on the word training.

Using our mediated models for those assigned to learn <u>control words</u>, control word usage directly increased satisfaction with life, with no potential mediating effects of either self-reported or behaviorally-derived EG. These results are not fully consistent with the set of Lasso models that explored this outcome. In those analyses, we did find that control word usage interacted with TMMS_MR scores in their relationship to satisfaction of life. The Lasso models showed some additional effects for control words on additional outcomes. Specifically, control word accuracy showed a simple effect on emotion dysregulation when not allowed to interact with other variables, but both control word usage and accuracy also interacted with various aspects of self-reported EG measures when allowed (TMMS_AF and TMMS_CF) to affect dysregulation. More interestingly, control word understanding interacted with the behavioral measures of EG (ICC_Neg) to affect emotional dysregulation, which was a notable difference from those trained on emotion words. Additionally, control word usage interacted with the TMMS_MR to affect satisfaction with life. Finally, the Lasso models showed that control word understanding and usage had simple effects on mental health, which differed from emotion word knowledge variables.

### Alignment with Previous Findings

The current findings generally align with prior research emphasizing the role of emotional vocabulary (and to some extent, any vocabulary) in enhancing emotion regulation and well-being [1, 7–8, 50]. The planned analyses showed little support for EG mediating the relationship between emotion word knowledge and outcomes, with most effects being directly between predictors and outcomes, and/or between mediators and outcomes. That said, individuals assigned to learn emotion words (although they surreptitiously also increased some aspects of control word learning) showed increases in EG measures and/or predicted decreases in suppression and emotional dysregulation. These findings are consistent with studies that show enhancing EG reduces emotional dysregulation and improves adaptive emotional strategy use [10–11], and they are consistent with the idea that EG increases subtle distinctions between emotions that, in turn, enable effective regulation strategies [16]. They also reinforce the finding that EG is associated with emotional clarity [52–53], and possibly awareness (using the TMMS_AF scale) [54], and that differentiation and range of emotional experiences are all related aspects of EG. Finally, they support the idea that emotion word knowledge (primarily understanding and accuracy) reduces emotion dysregulation beyond self-reported EG scores [50]. It is worth noting, however, that individuals assigned to learn control words (who also surreptitiously improved some aspects of emotion word learning while reducing other aspects of emotion word learning) showed improvements in satisfaction with life without affecting EG measures.

While prior research has emphasized the protective effects of high EG (compared to low EG) [26, 28], the current findings suggest that increasing emotion vocabulary might have similar effects. Specifically, increased usage of emotion words might reduce emotional dysregulation through participants’ *beliefs* about their EG abilities (i.e., the self-reported EG measures). The lack of an observed effect of the behaviorally-derived EG (ICC_Neg) in this study contrasts with previous work [45], however. This discrepancy may be attributed to how our study calculated the average ICC from negative emotions over the entirety of training, which did not capture momentary changes in variability in real-world emotional experiences. To this end, we did not find that self-reported surveys of EG and behaviorally-derived estimates of EG were correlated. This disconnect highlights potential differences in what these instruments capture. ICC scores may measure situational variability in emotional labeling, while self-reported measures may better capture trait-like emotional differentiation, or even *beliefs* about one’s ability. Therefore, ICC-based EG might be more context-sensitive and less reflective of enduring emotional skills. The relationship between ICC scores and self-reported measures warrants further investigation to clarify their respective contributions to the construct of EG, and to determine whether EG is a unified construct or rather is comprised of individual skills of attention, emotion clarity, and emotional differentiation.

### Limitations

Although we obtained our desired sample size based on medium indirect effects, the study might have been underpowered to detect smaller mediation effects or higher-order interactions. Additionally, the emotion word set used in the intervention was necessarily limited in scope; broader training inventories may yield stronger or more generalizable effects. Although the 14 granular emotion words used in the emotion word training protocol were well piloted and normed, they represent only a handful of the possible emotion words that could be used as training words.

Another limitation is that the majority of participants self-identified as highly educated, female, and Caucasian, and this study was conducted in one metropolitan city in the United States. This limits the generalizability of these results to individuals of other ethnicities, countries, education levels, and genders.

### Future Directions

The current study paves the way for future therapeutic interventions. The improvements observed in the emotion word group (and to a differing extent in the control word group) suggest that training improves emotional dysregulation and limits the use of emotional suppression. Future research should explore whether and how suppression is associated with reappraisal under various conditions [71]. Furthermore, this research shows that interventions can be cost-effective and easily executed, further expanding the breadth of clinical applications. Future research should analyze the effects of an emotion word training intervention on clinical populations that struggle with emotion regulation, such as those with mood disorders or anxiety disorders. Consequently, interventions targeting emotion vocabulary may yield especially strong benefits for clinical populations characterized by difficulties identifying and differentiating emotional experiences.

Additionally, future studies should examine whether similar interventions might be used to aid in effective emotion development. This might include introducing emotion vocabulary training earlier in development, which produces long-term benefits for emotional competence, resilience, and mental health. School-based interventions may represent a particularly promising context for cultivating EG before the onset of significant psychopathology.

Finally, future studies should investigate whether different operationalizations of EG provide a more nuanced understanding of the construct. This would include investigating how other measures relate to the nomological net of skills under the umbrella of “granularity”. Additionally, because emotion concepts are shaped by language and culture [8], cross-cultural research is needed to determine whether the effects of emotion word training generalize across linguistic and cultural contexts.

In summary, by demonstrating that interventions targeting emotion word knowledge can improve emotion regulation and well-being, this study underscores the practical applications of word-training research. Longitudinal studies might examine the durability of these effects and whether they translate into reduced vulnerability to psychopathologies over time. Future work should examine whether baseline granularity, symptom severity, developmental stage, or cognitive and linguistic abilities moderate intervention outcomes. Such findings could inform the development of more personalized emotion-focused treatments.

## Acknowledgements

The authors would like to thank members of the lab, including Charlie Adams, Sarah Ahmad, Hiral Amin, Ryan Anderson, Payton Chance, Leigh Cox, Sana Ismail, Savannah Kiser, Yvette McRoy, Payal Morari, Rochely Negron, Reza Razvi, David Ting, and Divya Venkat, who assisted with stimuli development, data coding, and the programming and running of participants.

## Competing Interests

The authors declare no competing interests.

## Funding

Funding was obtained from [(University removed for identity)] through a Provost’s grant awarded to the first author in 2023-2024, entitled “The Effects of Teaching Emotional Granularity through Emotion Words on Well-Being, Emotional Regulation, and Emotion Perception”.

## Disclosures

The authors have no disclosures, competing interests, or conflicts of interest. The manuscript’s main text contains an explicit statement that all studies, measures, manipulations, and data/participant exclusions are reported in the manuscript. The study was approved IRB 1906743 on September 6, 2022. Written consent was obtained. Data collection began September 2022 and continued August 2024.

## Data Availability

Data may be accessed by emailing the first author or viewed on https://osf.io/3ephu/overview?view_only=fe5d7dea2be643d4ba9046f31c724587. This study was not pre-registered.

## Author Contributions

First author conceived of the idea, developed the stimuli, aided in the collection of data, and performed all analyses with assistance. She wrote the majority of the manuscript. Second Author assisted with stimuli development, software programming, and data analysis. He assisted in the writing of the manuscript. Third Author assisted with data analysis and figure formatting. She assisted in the writing of the manuscript and approved the final edits. Fourth and Sixth Authors assisted with data analysis and assisted in the writing of the manuscript. Fifth Author performed Lasso model and assisted in the writing of the final manuscript.

## Notes

### Competing Interest Statement

The authors have declared no competing interest.

